# Reprogramming protein kinase substrate specificity through synthetic mutations

**DOI:** 10.1101/091892

**Authors:** Joshua M. Lubner, George M. Church, Michael F. Chou, Daniel Schwartz

## Abstract

Protein kinase specificity is largely imparted through substrate binding pocket motifs. Missense mutations in these regions are frequently associated with human disease, and in some cases can alter substrate specificity. However, current efforts at decoding the influence of mutations on substrate specificity have been focused on disease-associated mutations. Here, we adapted the Proteomic Peptide Library (ProPeL) approach for determining kinase specificity to the task of exploring structure-function relationships in kinase specificity by interrogating the effects of synthetic mutation. We established a specificity model for the wild-type DYRK1A kinase with unprecedented resolution. Using existing crystallographic and sequence homology data, we rationally designed mutations that precisely reprogrammed the DYRK1A kinase at the P+1 position to mimic the substrate preferences of a related kinase, CK II. This study illustrates a new synthetic biological approach to reprogram kinase specificity by design, and a powerful new paradigm to investigate structure-function relationships underpinning kinase substrate specificity.

## INTRODUCTION

Through their role in the covalent transfer of phosphate from a donor ATP molecule to a phosphoacceptor serine, threonine or tyrosine in a substrate protein, protein kinases in eukaryotes play key roles in cellular signal transduction, and function as gatekeepers for important events such as cell cycle checkpoints, apoptosis, and the immune response (1, 2). There are several levels of specificity that allow an individual protein kinase to navigate the daunting number of potential substrates, target the correct subset of proteins, and the correct residues within the appropriate protein for phosphorylation. Beyond temporal and spatial colo-calization, protein kinases also attain substrate specificity through pattern recognition of distinctive residues proximal to the phosphoacceptor residue (the “P-site”). This pattern is referred to as a kinase specificity motif (or simply “motif”), and is a model of substrates that are compatible with the kinase’s substrate binding pocket and can thus be phosphorylated. Motifs are primarily inferred from known physiological substrates (3), and are sometimes modeled as a string of allowable residues, as a position weight matrix, or as a combination of these. The presumed motif is a well-established starting point for *in silico* prediction of putative substrates (4); however, for nearly all protein kinases, the numbers of known substrates are very few in number resulting in poorly defined, low-resolution motif models.

Recently, Creixell and colleagues demonstrated several cancer mutations within kinase domains that modulated catalytic activity, and in some cases altered substrate specificity (5). These results, along with previous work that traced evolutionary changes in substrate specificity to amino acid substitutions (6), and the identification of potential specificity-determining positions (7, 8), suggest that a thorough investigation of amino acid structure-function relationships will be necessary to achieve a principled understanding of kinase specificity. At present, these studies have largely evaluated individual, naturally occurring kinase mutations. In this work, we sought to explore the potential for rational reprogramming of kinase substrate specificity through multiple directed synthetic mutations.

Here, we used the <u>Pro</u>teomic <u>Pe</u>ptide <u>L</u>ibrary (ProPeL) method (9) to accurately measure the specific motifs of both wild-type and mutated kinases (Fig. 1). Using this approach, first a heterologous kinase of interest is expressed in *E. coli.* The kinase phosphorylates bacterial proteins consistent with its endogenous kinase specificity motif. The extremely low activity of serine/threonine/tyrosine kinases and phosphatases in *E. coli* (10) allows for a high signal-to-noise ratio, and the absence of confounding human kinase cascades ensures a direct link between expressed kinase and observed phosphorylation event. After cell lysis and proteolysis, the resulting phosphopeptides are identified by tandem mass spectrometry. This can provide hundreds to thousands of kinase-specific phosphopeptides from which a high-resolution motif model is generated. In this case, the motif model is a position weight matrix with constant residues at one or more positions, which are easily visualized using the pLogo (11) graphical representation. That these bacterial substrates are not physiological is irrelevant - the identified motif can be used to accurately model kinase substrate specificity, and predict human substrates (9). Here, we have repeatedly utilized ProPeL to generate and compare motifs for wildtype and synthetic mutant kinases.

**Fig. 1.**
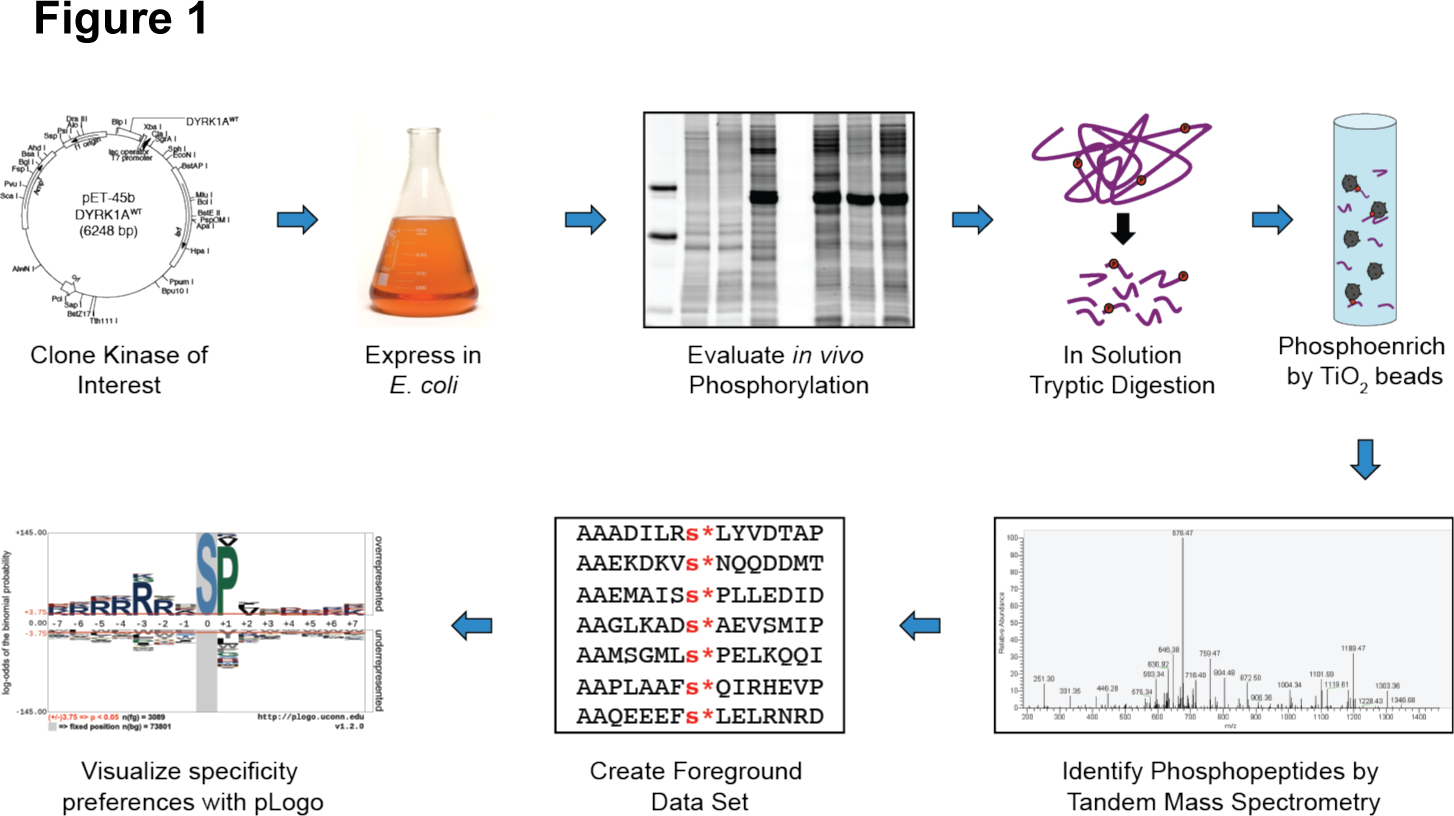
Schematic overview of ProPeL. A kinase of interest is cloned, and expressed in *E. coli.* Resulting bacterial phosphorylation is evaluated by SDS-PAGE with Pro-Q Diamond and Coomassie staining. Lysate is digested, phosphoenriched and identified by tandem mass spectrometry. Data sets are computationally analyzed with *motif-x (19)* and visualized with pLogo (11).

We chose the Down’s syndrome associated Dual specificity tyrosine-phosphorylation-regulated kinase 1A (DYRK1A) to act as a model kinase. Although the number of known human DYRK1A substrates is low (only 31, (12)), the specificity motif for wild-type DYRK1A has been partially characterized as including basophilic determinants, and a preference in the P+1 position for proline (13, 14), where P+n denotes the nth residue towards the C-terminus of the phosphoacceptor P-site, and P-n denotes the nth residue towards the N-terminus. Mechanistically, the region of the substrate binding pocket spanning the conserved DFG and APE residues within the kinase sub-domains VII - VIII is termed the “activation segment”, and has been implicated through X-ray crystallography to confer substrate specificity by interacting with the amino acids flanking the substrate’s P-site (reviewed in Kannan and Neuwald, 2004, (15)). DYRK1A is a member of the CMGC (<u>C</u>DK/<u>M</u>APK/<u>G</u>SK3/<u>C</u>LK) kinase family, and it has been suggested that the P+1 proline specificity typical of this family is imparted by a hydrogen bond with a CMGC-conserved arginine in the activation segment (Fig. 2A, (15, 16)). Given this model, we hypothesized that disrupting this hydrogen bond would reduce DYRK1A’s preference for proline at the substrate’s P+1 position. An interesting exception to the CMGC family P+1 proline preference is Casein kinase II (CK II), which prefers acidic residues at position P+1 (Fig. S1, (9, 17)). At the CMGC-conserved arginine position (R328 in DYRK1A), CK II instead codes for lysine (residue K198, (15)). Therefore, we predicted that the mutant DYRK1A^R328K^ (mimicking CK II at the CMGC arginine position) would re-position the lysine side-chain ε-amino group to allow for an electrostatic interaction with substrates containing a P+1 acidic residue.

**Fig. 2.**
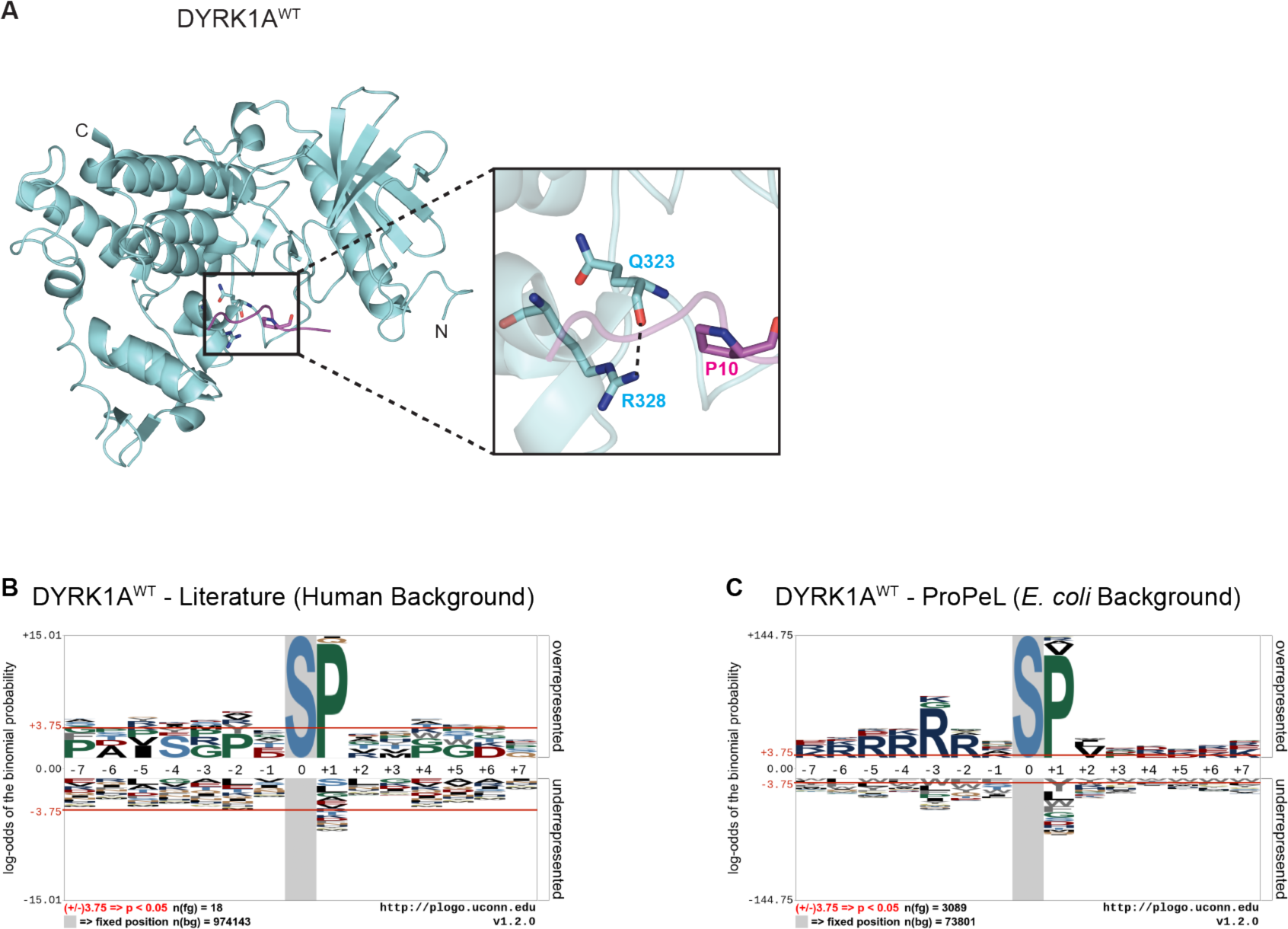
Crystal structure and high-resolution pLogos for DYRK1A^WT^. (**A**) Structural visualization (PDB Model 2WO6 (14)) indicating putative hydrogen bond between the main-chain oxygen of Q323 and a side chain nitrogen of R328 in the DYRK1A^WT^ activation segment. DYRK1A is colored in cyan, with peptide substrate in magenta. (**B** to **C**) pLogos (11) illustrate substrate preferences for DYRK1A^WT^, constructed from either (**B**) known literature-curated substrates (12) or (**C**) from unbiased ProPeL experiments. Overrepresented residues are displayed above the x-axis, underrepresented residues are below the x-axis. The *n(fg)* and *n(bg)* values at the bottom left of the pLogo indicate the number of aligned foreground and background sequences used to generate the image, respectively. The red horizontal bars correspond to p = 0.05 (corrected for multiple hypothesis testing), and y-axis is logarithmic scale. The grey box indicates a "fixed” residue. Additional pLogos for threonine-centered substrates are in Fig. S4.

In this work, we demonstrate the ability to generate kinase specificity models of unprecedented resolution using the ProPeL method. Using existing structural data and sequence homology, we successfully engineered the DYRK1A kinase to exhibit an unnatural substrate specificity, using both individual and multiple directed mutations. Overall, this study illustrates the effects of synthetic activation segment mutations upon substrate specificity, and introduces a new approach for the rational creation of designer kinases.

## RESULTS

### High-resolution determination of wild-type DYRK1A substrate specificity

Before attempting to reprogram DYRK1A, we first needed to create a sufficiently high-resolution model of wild-type DYRK1A substrate specificity to serve as a reference. We created bacterial expression constructs to express (1) a catalytic domain truncation of human DYRK1A (N137 - S496, referred to herein as DYRK1A^WT^); and (2) a catalytically inactive DYRK1A triple mutant K188R/D287N/D307N to function as a kinase dead negative control (referred to herein as DYRK1A^KD^). Both kinases express robustly in the C41(DE3) *E.coli* strain as evaluated by western blotting (Fig. S2, lanes 1-3). Using the in-gel phosphoprotein stain Pro-Q Diamond, we observed DYRK1A^WT^ exhibits strong autophosphorylation, and efficiently phosphorylate *E. coli* proteins throughout the gel and thus across the proteome. Importantly, DYRK1A^KD^ shows neither autophosphorylation, nor substrate phoshorylation (Fig. S3A and S3B, lanes 1-3).

**Fig. 3.**
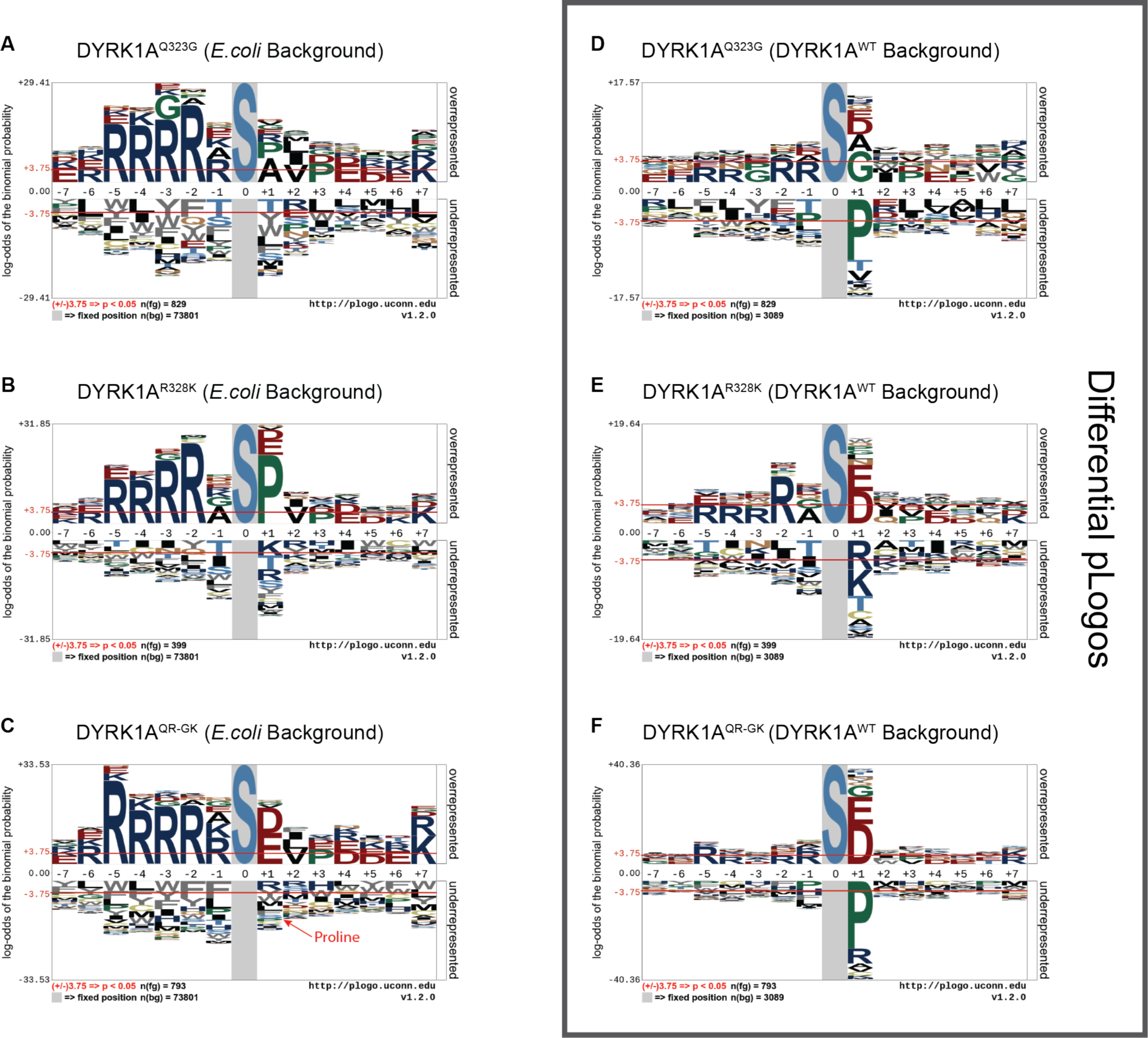
DYRK1A^QR-Gk^ exhibits altered P+1 substrate specificity. (**A** to **C**) pLogos (as described in Fig. 2) illustrate substrate preferences for (**A**) DYRK1A^Q323G^, (**B**) DYRK1A^R328K^, and (**C**) DYRK1A^QR-GK^ (**D** to **F**) Differential pLogos display the relative changes in substrate specificity between DYRK1A mutants and DYRK1A^WT^ by using DYRK1A^WT^ as a background data set, and mutant foreground data sets were used for (**D**) DYRK1A^Q323G^, (**E**) DYRK1A^R328K^, and (**F**) DYRK1A^QR-GK^ Additional pLogos for threonine-centered substrates in Fig. S7.

Using the ProPeL method, we identified 6,059 unique DYRK1A phosphorylation sites (3,089 pSer, 2,412 pThr, and 558 pTyr) on bacterial proteins. Note that this data set is an order of magnitude larger than that of the human kinase with the largest number of known natural substrates (CDK2, with 514 substrates (12)). Therefore, DYRK1A^WT^ ProPeL data results in high-resolution motifs that offer a dramatic improvement over those created with only the 18 known serine and 13 known threonine literature sites (visualized with the pLogo tool in Fig. 2B, 2C and Fig. S4A, S4B). In agreement with previous studies (13, 14), DYRK1A^WT^ exhibited a strong preference for proline in the P+1 position, and basic residues (particularly arginine) in the upstream positions, which together form the optimal consensus sequence RxxS*P (Fig. 2C). Our data confirms the recent suggestion in the literature that DYRK1A can phosphorylate substrates with alternative residues in the P+1 position (14); beyond a strong P+1 preference for proline, DYRK1A^WT^ also efficiently phosphorylates substrates with hydrophobic residues (particularly valine and alanine), or arginine in the P+1 position. While these residues are statistically significant in the P+1 position for serine P-sites, they fail to occur at statistical significance for threonine P-sites (Fig. S4B). Threonine P-site substrate specificity, therefore, may be more dependent on the P+1 proline than are serine P-site substrates. There is also a strong, previously unreported hydrophobic cluster present at P+2 for both serine and threonine substrates. DYRK1A does not exhibit a significant phosphotyrosine motif (Fig. S4C), however our 558 unique tyrosine phosphorylation sites in *E. coli* clearly indicate that DYRK1A is capable of phosphorylating tyrosine substrates *in trans*, and that phosphotyrosine activity is not restricted to autophosphorylation, as previously thought (13, 18).

Using the pLogo tool and an internal version of the *motif-x* program (19, 20), we evaluated dependence between motif positions. It is important to note that while we identify tryptic peptides by tandem mass spectrometry, the kinase-substrate interaction that produced the phosphorylation event occurred in the context of full-length substrate proteins. Therefore, we are able to map tryptic fragments back to the known *E. coli* proteome and extend sequences beyond the detected tryptic fragment, allowing us to analyze the presence (or absence) of multiple upstream basic residues. This analysis revealed that although there is a strong correlation between P+1 proline and upstream basic residues (Fig. S5A, S5B), there is no significant correlation between multiple upstream basic residues (Fig. S5C to S5F). Therefore, the optimal motif sequence is actually RxxS*P, and not RRRRxS*P, which is the broad motif without respect to any interdependent substrate residues. Substrates conforming to RxS*P, RxxxS*P, and RxxxxS*P with single arginines are thus also favored, but less so than those with RxxS*P. We note that multiple arginines are actually *not* favored for substrate recognition, although they do not appear to be clearly disfavored either. The complete list of statistically significant motif classes for DYRK1A (and all kinases within this study) identified by *motif-x* can be found in Table S2.

To verify the specificity of our DYRK1A model amongst other known kinases, we performed an *in silico* analysis using our high-resolution DYRK1A motif. We scored known human DYRK1A substrates, an equivalent number of substrates randomly selected from the human proteome, as well as known substrates for *other* kinases from the remaining serine/threonine kinase families (12). Our motif was able to accurately discriminate known DYRK1A substrates from random substrates and also performed well in discriminating against non-DYRK1A kinase substrates (Fig. S6).

### Mutation of Q323 reduces wild-type DYRK1A P+1 proline preference

As introduced earlier, DYRK1A P+1 substrate preference is hypothesized to be imparted by a hydrogen bond between the side-chain nitrogen of the CMGC-conserved arginine (DYRK1A^R328^) and the main-chain oxygen of a non-glycine residue (DYRK1A^Q323^) undergoing torsional strain (Fig. 2A, (15, 16)). This hydrogen bond should thus neutralize the main-chain oxygen’s dipole moment and facilitate interaction with substrates containing a proline at P+1. The loss of the bulky glutamine side-chain in a glutamine to glycine mutant (DYRK1A^Q323G^) could theoretically relieve torsional strain and re-position the kinase’s main-chain oxygen. This could destabilize the hydrogen bond, and ultimately result in a reduced P+1 proline preference.

Using ProPeL to determine DYRK1A^Q323G^ specificity, we identified 1,579 unique phosphorylation sites (829 pSer, 678 pThr, and 72 pTyr). Consistent with our hypothesis, motif visualization with pLogo revealed a significant reduction in P+1 proline preference (Fig. 3A and Fig. S7A). For DYRK1A^WT^, hydrophobic residues constitute secondary determinants in the P+1 position (particularly for serine substrates). For our engineered DYRK1A^Q323G^ mutant acting upon serine P-sites, alanine, not proline, becomes the most statistically significant substrate residue at position P+1. For threonine P-sites, proline is still the most statistically significant residue at position P+1, but is greatly reduced when compared to DYRK1A^WT^.

### Mutation of R328 mimicks CK II P+1 acidic preference

As discussed above, we hypothesized that creation of a DYRK1A variant that mimics CK II at the CMGC-conserved arginine position (DYRK1A^R328K^) would reprogram the P+1 preference from proline to acidic residues. Using ProPeL to analyze DYRK1A^R328K^, we identified 756 unique phosphorylation sites (399 pSer, 327 pThr, and 30 pTyr). Agreeing with our hypothesis, the DYRK1A^R328K^ mutant does show a significant increase in acidic P+1 preference for both serine and threonine P-site substrates (Fig. 3B and Fig. S7B). We note that phosphorylation of substrates with P+1 proline occur at similar levels to DYRK1A^WT^, suggesting that the substitution of arginine with lysine at the CMGC-conserved arginine position is still capable of forming the putative hydrogen bond. In the absence of a crystal structure, one possible explanation is that the positively charged side-chain ε-amino group of DYRK1A^R328K^ has been repositioned, and now facilitates a favorable interaction with substrates possessing an acidic residue at position P+1.

### Double mutation of residues Q323 and R328 completely reprograms DYRK1A P+1 specificity

Given the potentially independent influence of each of the DYRK1A^Q323G^ and DYRK1A^R328K^ mutations on the substrate P+1 specificity, we decided to investigate if the double mutant (referred to herein as DYRK1A^QR-GK^) would more fully recapitulate CK II specificity at P+1 than either individual mutant. In total, DYRK1A^GR-GK^ ProPeL experiments led to the identification of 1,535 unique phosphorylation sites (793 pSer, 645 pThr, and 97 pTyr). Together, the two mutations exhibited a combined effect on specificity, resulting in the complete reprogramming of the P+1 position from proline (characteristic of DYRKIA^WT^) to acidic residue preferences (mimicking CK II, Fig. 3C and Fig. S7C). Indeed, proline completely shifted from being the most dominant feature of DYRKIA^WT^ at position P+1 to statistical underrepresentation in DYRK1A^QR-GK^. Interestingly, the effect of the double mutant on substrate preference appears largely localized to the P+1 position, and maintains the upstream basic as well as the P+2 hydrophobic preferences of DYRK1A^WT^.

### *In silico* differential analysis offers additional insight

Although differences between the kinase specificity motifs for DYRK1A variants can be observed by simple inspection of the respective pLogos, it can be difficult to reconcile relative deviations from the wild-type kinase, especially when the sizes of the foreground datasets vary. To facilitate the direct comparison of each mutant DYRK1A with the wild-type kinase, we performed an *in silico* differential analysis between the DYRK1A^WT^ and mutant DYRK1A data sets. Typically pLogos will highlight significant motifs in kinase substrates by using phosphorylation sites as a foreground data set, and amino acid frequencies from the organism’s proteome to determine background probabilities. In order to directly compare each DYRK1A mutant to DYRK1A^WT^, we still used each mutant DYRK1A data set as the foreground, but then used our 6,059 DYRK1A^WT^ phosphorylation sites instead of the *E. coli* proteome as a background data set. The resulting “differential pLogos” display residues that are over- and underrepresented in the respective mutant DYRK1A substrate pool *relative* to the wild-type kinase (DYRK1A^WT^) substrates, rather than the background proteome.

As already noted, DYRK1A^Q323G^ differs from wild-type by favoring proline less than DYRK1A^WT^ at P+1 and shifting P+1 preference to alanine, and in the differential pLogo, that shift is abundantly clear (Fig. 3D and Fig. S7D). Next, the differential pLogo for DYRK1A^R328K^ (Fig. 3E and Fig. S7E) indicates no shift to disfavor proline at P+1 relative to DYRK1A^WT^, which further confirms the standard pLogo results. A striking feature of the differential pLogo (that is less obvious in the standard pLogo) is an overrepresentation of acidic residues with a concomitant underrepresentation of basic residues at P+1 compared to DYRK1A^WT^. For DYRK1A^QR-GK^, the differential pLogo confirms the additive effects of the two mutants, namely the disfavoring of proline and the favoring of acidic residues at the P+1 position. We also observed that while arginine remains a positive determinant in the upstream positions for all DYRK1A variants (Fig. 3A to 3C, and Fig. S7A to S7C), the importance of this upstream region increases slightly but not uniformly in the mutants (note the more significant P-2 arginine in differential pLogos Fig. 3E and Fig. S7D, S7E, but not in Fig. 3D, Fig. 3F nor Fig. S7F). This suggests that in addition to reprogramming the P+1 substrate preference, there may be some subtle rearrangement of the kinase pocket, resulting in the observed shift in upstream basophilic preferences.

## DISCUSSION

Recently, catalytic domain kinase mutations associated with cancer have been linked to altered substrate specificity, and modulated catalytic activity (5). While there have been previous efforts at engineering kinases within the bacterial two-component system (21, 22), identification of important residues influencing phosphoacceptor preference (7, 8), and tracing evolutionary lineage through ancestral kinases (6), these studies have all exploited natural variation among kinases. In the present study, we rationally engineered DYRK1A to first abolish its endogenous P+1 proline preference (which interestingly most closely resembles the specificity of the unrelated kinase PKA). By incorporating a second point mutation to make our DYRK1A^QR-GK^ double mutant, we successfully shifted P+1 preference to favor acidic residues and disfavor proline. This combination results in a completely synthetic hybrid specificity that combines both upstream DYRK1A and CK II P+1 preferences.

In large part, this investigation was made possible by the unprecedented resolution achievable through the use of ProPeL, which provides an extremely detailed motif representation that extends far beyond what is available from known endogenous substrates. It is readily apparent that our approach can be scaled up to a more systematic interrogation of structure-function relationships of other residues within the activation segment. The observation that the DYRK1A^R328K^ mutant (CK II mimic) began to exhibit canonical CK II substrate preferences, while the DYRK1A^Q323G^ mutant reduced proline preference by a seemingly distinct mechanism, demonstrates that there is some independence of residues within the activation segment, and it may be possible to program other effects without crosstalk. Such an investigation would produce critical insights into the mechanistic underpinnings of kinase specificity and the residues relevant to kinase-substrate interactions.

Exploring the mutation space of the activation segment may also help define the sensitivity of protein kinases to mutational burden (23). As the number of disease-associated missense mutations that localize to the catalytic domain continue to rise (12, 24, 25), our approach provides a powerful system for identifying mutation-induced kinase specificity rewiring. Ultimately, such analyses may be invaluable for drug design and for cancer therapeutics that often result in drug-resistant somatic kinase mutations with unintended consequences.

## MATERIALS AND METHODS

### Plasmids, strains and *in vivo* proteome phosphorylation

A plasmid containing the full-length coding sequences for the human DYRK1A gene in the pDNR-Dual vector was purchased from the Harvard PlasmID Repository (Boston, MA). The catalytic domain for DYRK1A (N137 - S496) was cloned into the pET45b vector (Novagen) by traditional restriction site PCR cloning. All mutations were performed according to the Stratagene QuickChange II protocol. DYRK1A constructs were expressed in the *E. coli* OverExpress C41(DE3) strain (Lucigen) by IPTG induction. DYRK1A expression was optimal when induced at mid-log with 0.5 mM IPTG and incubated for 3 hours at 37°C with shaking at 250 rpm. Cells were harvested by centrifugation at 6,000 g, 4°C for 10 minutes, and stored at −80°C.

### Lysis and analysis of *in vivo* phosphorylation

Cell lysate was prepared as described previously (9, 26) with minor modifications. Cells were lysed by sonication with a Fisher Sonic Dismembrator F60 at 15% power using 15-20 second pulses, with 1 minute rest on ice between pulses, until lysate was clear. Crude lysate was clarified by centrifugation at 20,000 g and 4°C for 30 minutes. Protein concentrations were determined by Bichinchoninic Acid (BCA) Assay (Pierce), phosphorylation level was evaluated by SDS-PAGE with Pro-Q Diamond Phosphoprotein stain (Life Technologies), and total protein was evaluated by GelCode Blue coomassie staining (Life Technologies).

### Western Blotting

Western blotting for DYRK1A used the primary antibody Anti-6xHis (NeuroMab clone N144/14, RRID: AB_10671171, UC Davis/NIH NeuroMab Facility) at 1:1000 dilution, and IRDye 800CW Goat anti-Mouse IgG secondary antibody (LI-COR Biosciences) at 1:5000 dilution.

### In solution tryptic digestion

Samples were reduced, alkylated, digested with trypsin (Sequencing grade modified, Promega; bovine trypsin, Sigma; or TrypZean, Sigma) at a 1:100 enzyme:substrate ratio, and desalted with either 100 mg or 500 mg tC18 SepPak Vac solid-phase extraction cartridges (Waters) as previously described in Villén and Gygi, steps 2-17 (26). Desalted peptides were snap-frozen in liquid nitrogen and lyophilized. Peptides were resuspended in appropriate buffer for one or more of the following phosphopeptide enrichment strategies.

## PHOSPHOENRICHMENT

Over the course of many mass spectrometry runs, many different phosphoenrichment strategies were evaluated. Ultimately, we concluded that the most efficient sample preparation was a simple TiO_2_ enrichment step, as described below. However, data from the other methods were collected and accumulated for the DYRK1A^WT^ pLogo, and as such is summarized below. All mutant DYRK1A data were obtained using simple bulk TiO_2_ enrichment.

### TiO_2_ bead enrichment

Phosphopeptide enrichment using bulk TiO2 beads (Titansphere 5 μm, GL Sciences) was modified from Kettenbach and Gerber [16]. Beads were conditioned in bulk using Binding Buffer (50% ACN, 2 M Lactic Acid), with beads added at a 4:1 ratio to peptides (peptide concentration estimated by NanoDrop A280 absorbance), and brought to a final peptide concentration of 1 mg/mL. Peptide/bead mix was incubated with maximum shaking on an Eppendorf Thermomixer at room temperature for 1 hour. Beads were washed with Wash Buffer (50% ACN, 0.1% TFA) and eluted with 5% NH_4_OH. Eluate was immediately acidified by addition of FA, dried in a speed-vac, and stored at −20°C for further enrichment, or analysis by mass spectrometry.

### Strong Cation Exchange (SCX)

Traditional SCX by HPLC was performed as described previously (9, 26). Separation by SCX-SPE was performed according to Dephoure and Gygi (27). When SCX-SPE was performed prior to TiO_2_ phosphoenrichment, fractions were desalted using 100 mg tC18 SepPak Vac cartridges. For SCX-SPE performed subsequent to TiO_2_ phosphoenrichment, fractions were desalted using in-house StageTips (28) packed with 5 C18 discs per tip. Desalted samples were dried in a speed-vac and stored at −20°C for further enrichment, or analysis by mass spectrometry.

### Electrostatic Repulsion Hydrophilic Interaction Chromatography (ERLIC) by SPE

ERLIC-SPE followed a similar principle as SCX-SPE, using the bulk material from ERLIC SPE WAX Macrospin columns (The Nest Group). ERLIC-SPE columns were conditioned with successive washes with methanol followed by water. Columns were incubated in 0.2 M NaH_2_PO_4_, 0.3 M NaOAc for >1 hour. Columns were equilibrated with 70% ACN, 20 mM Na-MePO_3_ (pH 2.0), and samples were loaded and washed with this same buffer. Peptides were eluted sequentially first with 10% ACN, 20 mM Na-MePO_3_ (pH 2.0), followed by 50 mM NaH_2_PO_4_. Final elution was achieved with 300 mM NaH_2_PO_4_. Eluates were desalted with either SepPak Vac cartridges or StageTips depending on volume, dried in a speed-vac and stored at −20°C for either further enrichment, or analysis by mass spectrometry.

### Peptide Identification by Tandem Mass Spectrometry

Peptides were resuspended in 30 Buffer A (3% ACN, 0.125% FA) and 1 4 loaded onto a C18 nanocapillary column with a pulled tip that sprays directly into the inlet of a Thermo Fisher Scientific LTQ Orbitrap XL mass spectrometer. Peptides were eluted using an Agilent 1200 HPLC binary pump with a gradient that changes solvents from 100% to 65% Buffer A (0% to 35% Buffer B) over a 48, 85, or 145 minute time period, where Buffer A = 3% ACN, 0.125% FA in water, and Buffer B = 0.125% FA in ACN. A T0P10 method was used (MS scans followed by Collision Induced Dissociation MS/MS on the top 10 most intense MS spectral peaks). Spectra were searched using SEQUEST against the E. coli proteome, including decoy database entries, which allowed for differential serine, threonine, and tyrosine phosphate modifications (+79.966331), a differential methionine oxidation modification (15.9949146221) and a constant cysteine modification of +57.02146374. The deltaXCORR (the difference between the first and second hits to the databases) was set to be >= 0.08. To minimize false positives, for each of the two classes of peptide charges z = +2 and z >= +3, XCORR thresholds were chosen to accept peptides in such a manner that 1% of them were hits from the decoy database, resulting in an expected False Discovery Rate (FDR) of 2%.

### Phosphopeptide list filtering

Prior to motif analysis, a master negative control list was generated by pooling phosphopeptides previously identified in negative control experiments (9), previously identified endogenous *E. coli* phosphorylation sites (10, 29), and phosphorylation sites identified in empty vector and kinase dead negative control experiments. Phosphorylation sites on this master negative control list were removed from each active DYRK1A variant data set to generate a final list of kinase-specific phosphorylation sites. Peptide lists from all runs were merged within each kinase variant, and redundant peptides were removed prior to motif analysis.

### pLogo Generation

To generate graphical motifs, known as pLogos, we used the online tool at plogo.uconn.edu, previously described in detail (11). See supporting Information for a more detailed explanation, and instructions for recreating pLogos with our provided data.

### Scoring known kinase substrates

*scan-x* analyses of known and random substrates were carried out using an internal version of the *scan-x* software (30). Candidate peptides were scored for a goodness-of-fit using our DYRK1A^WT^ position weight matrix (PWM) obtained through ProPeL. Known verified human substrates were retrieved from the PhosphoSitePlus database (12) (http://phosphosite.org), while random substrates were obtained by randomly choosing an equivalent number of serine/threonine 15 mers from the human proteome. Note that any substrate which was unable to be extended to a P-site centered 15mer due to proximity to either the N-or C-terminus was unable to be scored.

### *motif-x* Analysis

*motif-x* analyses were carried out using an internal version of the *motif-x* web tool (19) with the following parameters selected: central residue = S*,T*, or Y*, width = 15, foreground occurrence threshold = 5, significance threshold = 0.00001, background database = NCBI *E. coli* proteome, and background central residue = S,T, or Y.

### Structural Modeling

All structural modeling was visualized using PyMol for Mac, and using PDB Model 2WO6, retrieved from the RCSB Protein Data Bank (14).

## SUPPLEMENTARY MATERIALS

Fig. S1. pLogos for CK II, curated from known literature sites.

Fig. S2. Western blot for all DYRK1A kinase variants.

Fig. S3. SDS-PAGE gels for all DYRK1A kinase variants, using Pro-Q Diamond staining for phosphorylation activity, and normalized to total protein Coomassie staining.

Fig. S4. Additional pLogos for DYRK1A^WT^, including pLogos for threonine-and tyrosine-centered substrates (curated from known literature sites, and ProPeL experiments, and pLogos for a smaller subset of the data.

Fig. S5. Additional pLogos for DYRK1A^WT^ with different positions "fixed” to show conditional probabilities, and demonstrate multiple position correlations.

Fig. S6. Average position weight matrix (PWM) scores using the DYRK1A^WT^ pLogo to score substrates of DYRK1A, non-CMGC family kinases, or random phosphoacceptors, curated from the literature.

Fig, S7. Additional threonine-centered pLogos for mutant DYRK1A kinase variants.

Table S1. Mass spectrometry data.

Table S2. *motif-x* runs

Table S3. Aligned data

## Acknowledgements

The authors wish to thank Noah Dephoure and Craig Braun in the Gygi Lab for their assistance and support with mass spectrometry experiments. We thank Sean Lubner, John Redden, Stephanie Reeve, Megha Sah, Ben Stranges, and Randall Walikonis for generously sharing reagents and/or providing invaluable technical advice. We also thank Prakhar Bansal for maintainance and assistance with the Schwartz Lab computational tools. We thank Sean Congdon, Lylah Deady, Fred Murphy, Benjamin Currall and Anastasios Tzingounis for helpful discussions and the critical reading of our manuscript. Finally, we thank the University of Connecticut Computational Biology Core for hosting the pLogo web site, and maintaining the clusters on which it runs.

## Author Contributions

J.M.L. and D.S. conceived of the study. J.M.L., M.F.C., and D.S. designed the experiments. J.M.L. and M.F.C. performed the experiments and analyzed the data. G.M.C. and D.S. contributed materials, resources, and analysis tools. J.M.L. wrote the manuscript. All authors helped edit the final manuscript.

